# Outer membrane tube formation by *Francisella novicida* involves extensive envelope modifications and is linked with type VI secretion and alterations to the host phagosomal membrane

**DOI:** 10.1101/2024.12.17.628953

**Authors:** Maheen Rashid, Shoichi Tachiyama, Shiwei Zhu, Hang Zhao, William D. McCaig, Jingchuan Sun, Hulin Li, Jun Liu, David G. Thanassi

## Abstract

*Francisella tularensis* is a Gram-negative, intracellular pathogen that causes the zoonotic disease tularemia. Due to its ease of dissemination and high lethality, *F. tularensis* is classified as a Tier 1 Select Agent with potential for misuse as a bioweapon. The mechanisms by which *Francisella* replicates intracellularly and interacts with the host during infection are not well understood. *Francisella* produces spherical outer membrane vesicles (OMV) and novel tubular extensions of its cell surface that are also released extracellularly. These OMV and outer membrane tubes (OMT) contain *Francisella* virulence factors and are produced in response to amino acid starvation and during infection of macrophages. To investigate how the OMT are formed, we used cryogenic electron tomography to examine the model *Francisella* species, *F. novicida*, during *in vitro* culture and within the macrophage phagosome. OMT formation involved progressive alterations of the bacterial envelope, resulting in extensions of both the inner and outer membranes. A dynamic cytoplasmic structure was present at the base of the OMT that extended into the tubes during elongation, together with cytoplasmic material. OMT produced within the macrophage phagosome were associated with changes to the phagosomal membrane, suggesting a role in phagosomal escape. Consistent with this, using confocal microscopy, we observed colocalization of the *Francisella* type VI secretion system with the OMT, both within bacteria and in released tubular vesicles. These findings reveal the cellular transformations that occur during membrane tubulation by *Francisella* and provide insights into the function of membrane-derived structures during host-pathogen interactions.

## INTRODUCTION

*Francisella tularensis* is a pleiomorphic, Gram-negative, facultative intracellular bacterium and the causative agent of the zoonotic disease tularemia^1–4^. Tularemia manifests in various forms based on how it is contracted. Typically, an infection acquired through the dermal route, upon contact with an infected animal or through the bite of a tick or other vector, develops into ulceroglandular tularemia. These ulcers are mostly non-lethal and resolve within a few days^4^. Pneumonic tularemia is the more serious form of disease, in which inhaling even a very small dose of aerosolized bacteria leads to severe pneumonia, with a mortality rate as high as 60% if left untreated^5^. Due to its low infectious dose, ease of aerosolization, and potential for high morbidity and mortality, *F. tularensis* has been developed as a bioweapon and is categorized by the Centers for Disease Control and Prevention as a Tier 1 Select Agent^6^.

There are two subspecies of *F. tularensis* with clinical significance: subsp. *tularensis* (type A), which is highly virulent; and subsp. *holarctica* (type B), which is less virulent in humans compared to subsp. *tularensis*^6^. Another species within the *Francisella* genus*, F. novicida,* has limited virulence in healthy humans while maintaining its capability to cause severe disease in animal models. *F. novicida* infection of host cells and animals shares many similarities with *F. tularensis* infection, and *F. novicida* has served as an important experimental strain that can be handled under biosafety level 2 conditions^7^. The molecular basis for *Francisella*’s virulence and extreme infectivity remain poorly understood. This, together with the absence of a licensed and effective vaccine, make it important to understand mechanisms of *F. tularensis* virulence and intracellular pathogenesis to support the advancement of effective vaccines and therapeutic strategies against tularemia.

A central feature of *Francisella*’s virulence is its ability to evade and interfere with host innate immune responses to promote its proliferation within host cells^2,3,8,9^. *Francisella* infects and replicates within a variety of host cells, with macrophages being a preferred intracellular niche^6,10,11^. Following internalization into macrophages, *Francisella* is initially contained with the macrophage phagosome. The bacteria suppress maturation of the phagosome, avoid or interfere with intracellular signaling pathways, and resist host defense mechanisms such as antimicrobial peptides and reactive oxygen species^2,3,8,9^. Within as soon as 30-60 min after uptake, *Francisella* escapes from the phagosome and enters the host cell cytosol, where bacterial intracellular replication occurs. Extensive replication of the bacteria eventually activates host cell death pathways, resulting in bacterial release and spread to neighboring host cells for subsequent rounds of infection^10,12,13^. Critical to *Francisella’s* ability to escape the phagosome is a genomic locus termed the *Francisella* pathogenicity island (FPI)^10,12,14–16^. The FPI contains genes encoding a type VI secretion system (T6SS), which is utilized by many Gram-negative pathogens to deliver virulence factors to host cells and is essential for phagosomal escape by *Francisella* spp.^14,15,17,18^. *Francisella* spp. also encode a type I secretion system and type IV pilus assembly pathway, which are linked to effector secretion^19–21^. However, *Francisella* spp. lack the canonical type III and type IV secretion systems that are typically used by intracellular bacteria for effector delivery to host cells^7^. Given its high pathogenicity and ability to replicate intracellularly while suppressing host responses, it is likely that *Francisella* utilizes additional pathways for virulence factor secretion.

Previous work demonstrated the production of outer membrane vesicles (OMV) by *Francisella* spp.^22–25^. OMV are spherical structures released from the outer membrane (OM) of Gram-negative bacteria with a variety of functions, including biofilm formation, transport of cargo, acquisition of nutrients, response to envelope stress and hostile environments, and interaction with other bacteria and host cells^26–29^. For pathogenic bacteria, OMV have been shown to deliver effector proteins to host cells and modulate host immune responses, supporting a role for OMV as a virulence factor secretion system^26,29^. In addition to spherical OMV, *Francisella* spp. produce unique tubular vesicles and tubular extensions of its cell surface, termed outer membrane tubes (OMT)^23–25^. The OMV and OMT contain known *Francisella* virulence factors along with many hypothetical proteins^23–25^. Additionally, the OMV and OMT are produced during infection of macrophages and within the macrophage phagosome, suggesting they function during intracellular pathogenesis^10,23,30^. Mechanisms governing the regulation and production of bacterial membrane vesicles, and the selective packaging of vesicle contents, remain major questions in the field. In previous studies, we demonstrated that *F. novicida* produces OMV and OMT in a controlled manner that is regulated by specific genes and environmental signals^23,25^. We identified the inducing signal as amino acid starvation, including cysteine deprivation^25^. Amino acid starvation is experienced by the bacteria during infection of macrophages and is also a signal used by the major virulence regulator of *Francisella*, MglA, to turn on genes within the FPI, including the T6SS^31–33^. The OMT produced by *Francisella* are distinct from other bacterial tubular membrane extensions that have been described in the literature^34^ and mechanisms governing their production and function during infection are unknown.

In this study, we employed cryogenic electron tomography (cryo-ET) to conduct a detailed analysis of the *F. novicida* OMT. We observed a progressive alteration in bacterial morphology during tube formation, with dramatic changes occurring to the inner membrane (IM), periplasm, and OM. OMT formation occurred at the cell pole and involved extension of both the IM and OM. We detected a dynamic cytoplasmic structure that sits at the base of the tubes, adjacent to the IM. During OMT formation, the cytoplasmic structure changed shape and extended into the growing tubes, together with cytoplasmic material and the IM. Cryogenic electron microscopy (cryo-EM) imaging of OMT released from bacteria into the surrounding medium revealed that the released tubes also contained cytoplasmic and IM material. We used cryo-focused ion beam milling coupled with cryo-ET (cryo-FIB-ET) to visualize OMT produced by *F. novicida* inside the macrophage phagosome. Notably, alterations in the phagosomal membrane occurred at contact points between the OMT and phagosome, suggesting a role in phagosomal escape. Consistent with this, using confocal microscopy, we detected co-localization of the T6SS with OMT present on bacteria and released into the medium. These findings provide evidence for a sophisticated mechanism governing the generation of OMT and reveal their association with a key step in the intracellular pathogenesis of *Francisella*.

## RESULTS

### Cryo-ET imaging of *F. novicida* reveals dramatic changes to the bacterial envelope upon tube formation and the presence of novel cytoplasmic structures

We first used cryo-ET to examine *F. novicida* U112 grown under conditions in which the bacteria do not experience amino acid starvation and OMT production is not induced. The bacteria were grown for 44-48 h on brain heart infusion (BHI) agar supplemented with 0.1% (w/v) cysteine. *F. novicida* grown under these conditions exhibited the characteristic appearance of Gram-negative bacteria, with a relatively uniform, rodlike or coccobacillus shape (Fig. 1A and B, Movies S1 and S2). The bacteria contained a typical, dense cytoplasm populated with ribosomes, surrounded by a distinct IM, periplasm, and OM. The periplasm was mostly thin and evenly spaced, particularly along the long axis of the cell, as typical for Gram-negative bacteria. However, enlarged periplasmic regions were present, especially at the cell poles (Fig. 1A and B), and unevenness in the IM and OM was apparent, together with periplasmic structures of unknown identity that appeared to bridge between the IM and OM (Fig. 1C-F). Notably, distinct oval-shaped structures were observed within the cytoplasm of the bacteria, positioned adjacent to the IM and located at or near a cell pole (Fig. 1A, B, D and E, Movies S1 and 2). The oval cytoplasmic structures measured ∼50 to 125 nm in diameter and had a clearly defined border and lighter interior density than the surrounding cytoplasm. In some bacteria, two such cytoplasmic structures were present (Movie S2), and we also observed a more extended cytoplasmic structure in one bacterium (Fig. 1A and G).

**Figure 1.**
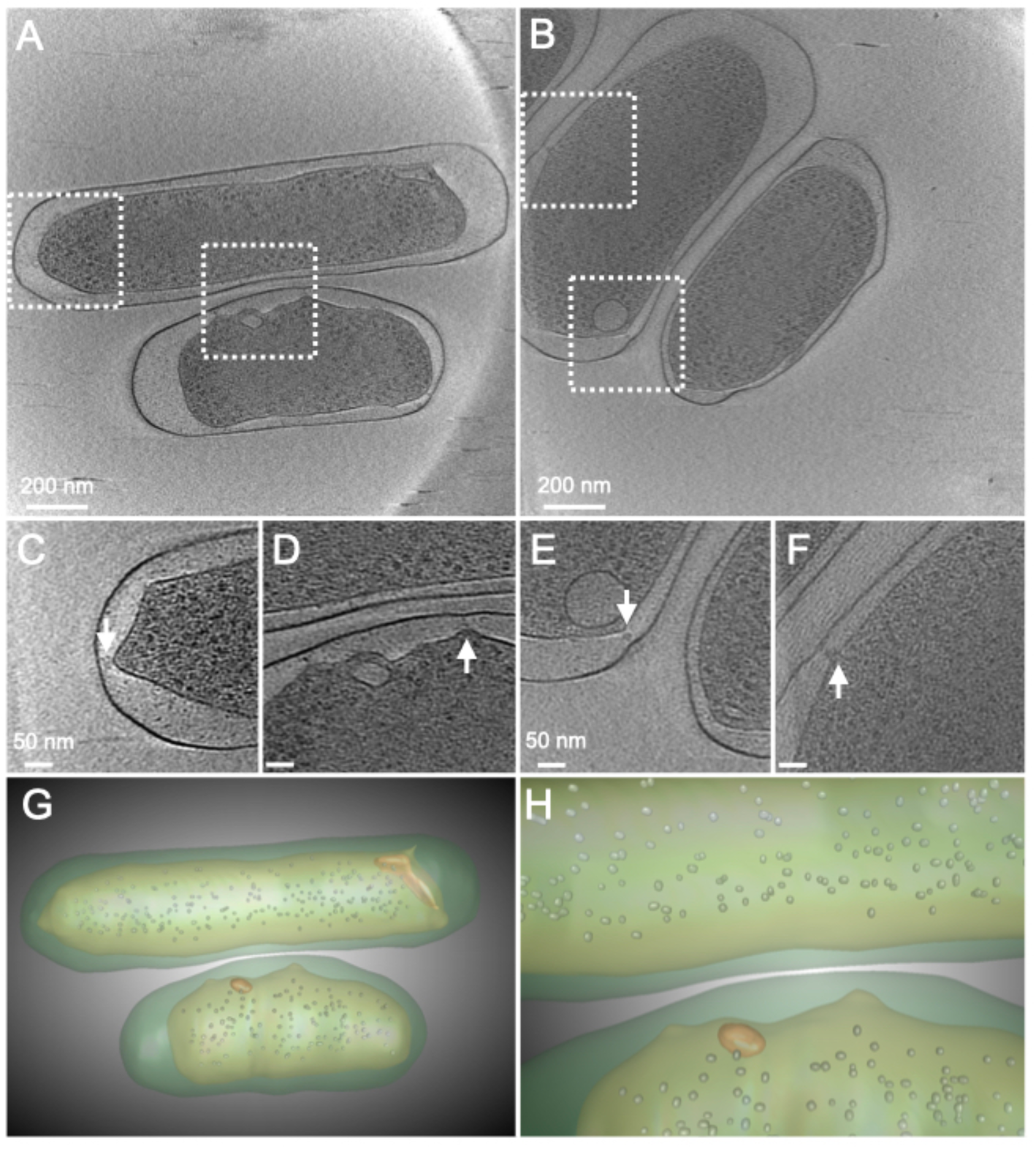
Cryo-ET imaging of *F. novicida* grown under cysteine sufficient (OMT repressing) conditions. (A and B) Single sections of tomograms showing *F. novicida* U112 grown on BHI agar supplemented with 0.1% cysteine supplementation. (C-F) Zoomed-in images of boxed regions in panels A and B, highlighting oval-shaped cytoplasmic structures and periplasmic structures that bridge the IM and OM (arrows). (G-H) Overview (G) and zoomed-in (H) segmentation images of the tomogram shown in panel A. The cytoplasmic structures are shown in orange and ribosomes (small round shapes) in gray. The cytoplasm and IM are indicated by light green, and the periplasm and OM by dark green.

We next examined *F. novicida* U112 grown for 44-48 h on BHI agar without cysteine supplementation. Under these conditions, the bacteria deplete the available cysteine, leading to amino acid starvation and the induction of OMT ^25^. Bacteria grown in these tube-inducing conditions exhibited substantial changes to their cell envelope and overall shape (Fig 2, Movies S3-5). The bacteria lost their rodlike morphology and became pleomorphic, with a more rounded cytoplasm and prominent bulges and distended regions of the periplasm and OM. Multiple extended regions of the OM could be observed per bacterium, typically at the cell poles (Fig. 2A-E). In some cases, one of these regions extended further to create a tubular projection from the bacterial cell surface (Fig. 2A, B, and D, Movies S3-5). Dense structures were visible in the cytoplasm of some bacteria (Fig. 2D and E), which may represent the formation of stress granules under the amino acid starvation conditions^35,36^. The previously observed oval cytoplasmic structures also underwent significant changes, enlarging and transforming into a more bulb-like conformation, the tip of which could be seen extending into the OMT projections (Fig. 2A-C, F and G, Movies S3-5). The bulb-like structure appeared to form the base or origin point of the OMT, with an IM tubule and associated cytoplasmic material extending further into the OM projection. In some cases, the cytoplasmic structure at the base of the OMT appeared to be fragmented (Fig. 2D and E, Movie S5). In these cases, the periplasm was less enlarged and the OMT was narrower, with condensed or aggregated material present inside the tube (Fig. 2D and E, Movie S5).

**Figure 2.**
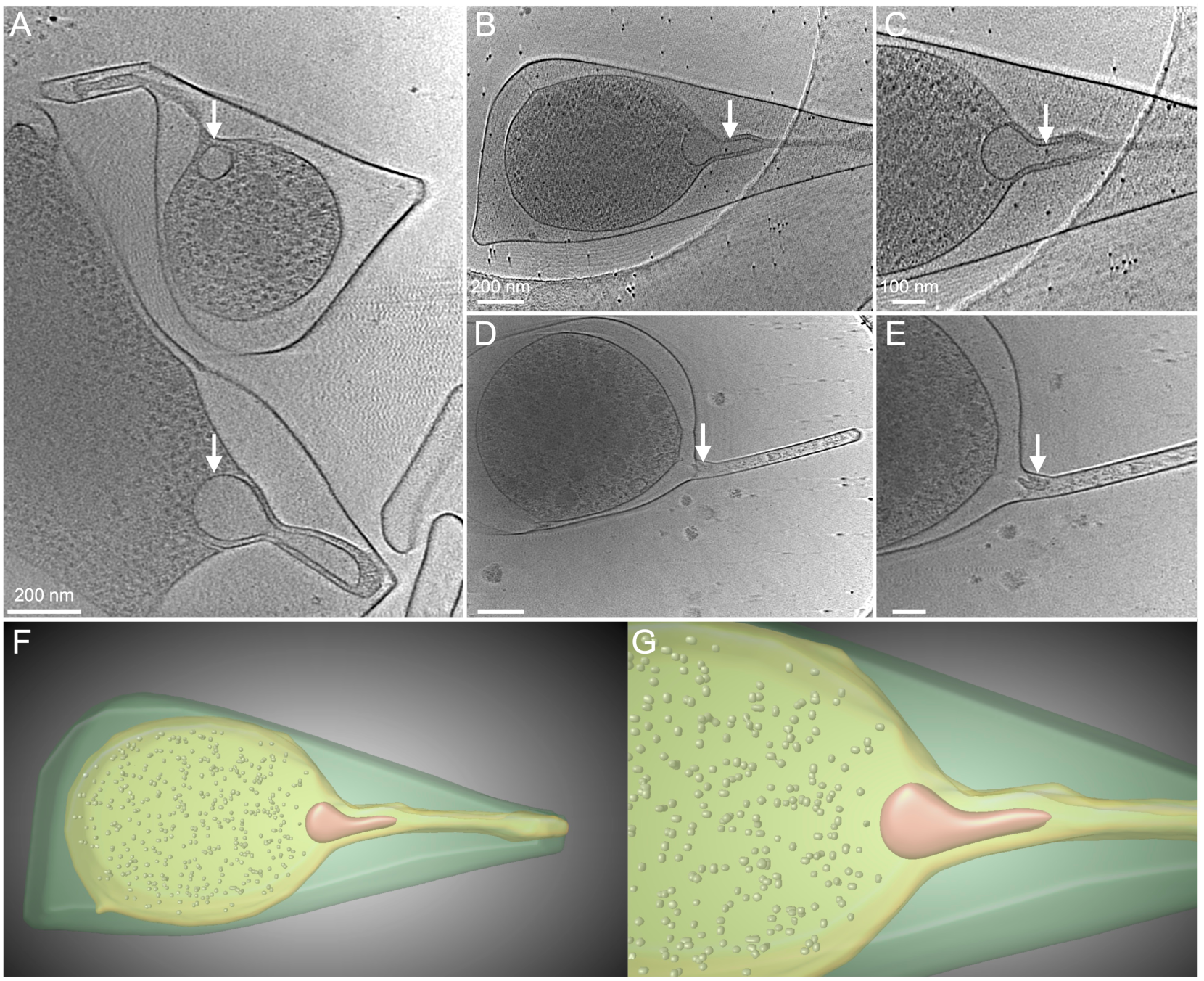
Cryo-ET imaging of *F. novicida* grown under OMT-inducing conditions. (A, B, and D) Single sections of tomograms showing *F. novicida* U112 grown on BHI agar without cysteine supplementation. (C and E) Zoomed-in images of panels B and D. Panes B and C show the cytoplasmic bulb-like structure (arrow) extending into the OM, along with cytoplasmic material. Panels D and E show a narrower OMT, with aggregated material present inside the tube (arrow) and a fragmented cytoplasmic structure. (F and G) Segmentation images of the tomogram corresponding to the images in panels B and C. The bulb-like structure and ribosomes are shown in orange and gray, respectively. The cytoplasm and IM are indicated by light green, and the periplasm and OM by dark green.

In addition to extending from the bacterial surface, the *Francisella* OMT are released into the surrounding medium, together with spherical OMV that are typical of Gram-negative bacteria^23–25^. We next used cryo-EM to examine vesicles released from *F. novicida* grown under tube-inducing conditions (BHI agar without cysteine supplementation). The vesicles exhibited a range of shapes and sizes, corresponding to both OMT and OMV (Fig. 3A). Internal structures were evident for many of the vesicles, with density and appearance consistent with cytoplasmic material surrounded by an IM (Fig. 3B-F). Thus, the cytoplasmic and IM regions observed extending into the tubes on the bacterial surface are maintained within the released OMT.

**Figure 3.**
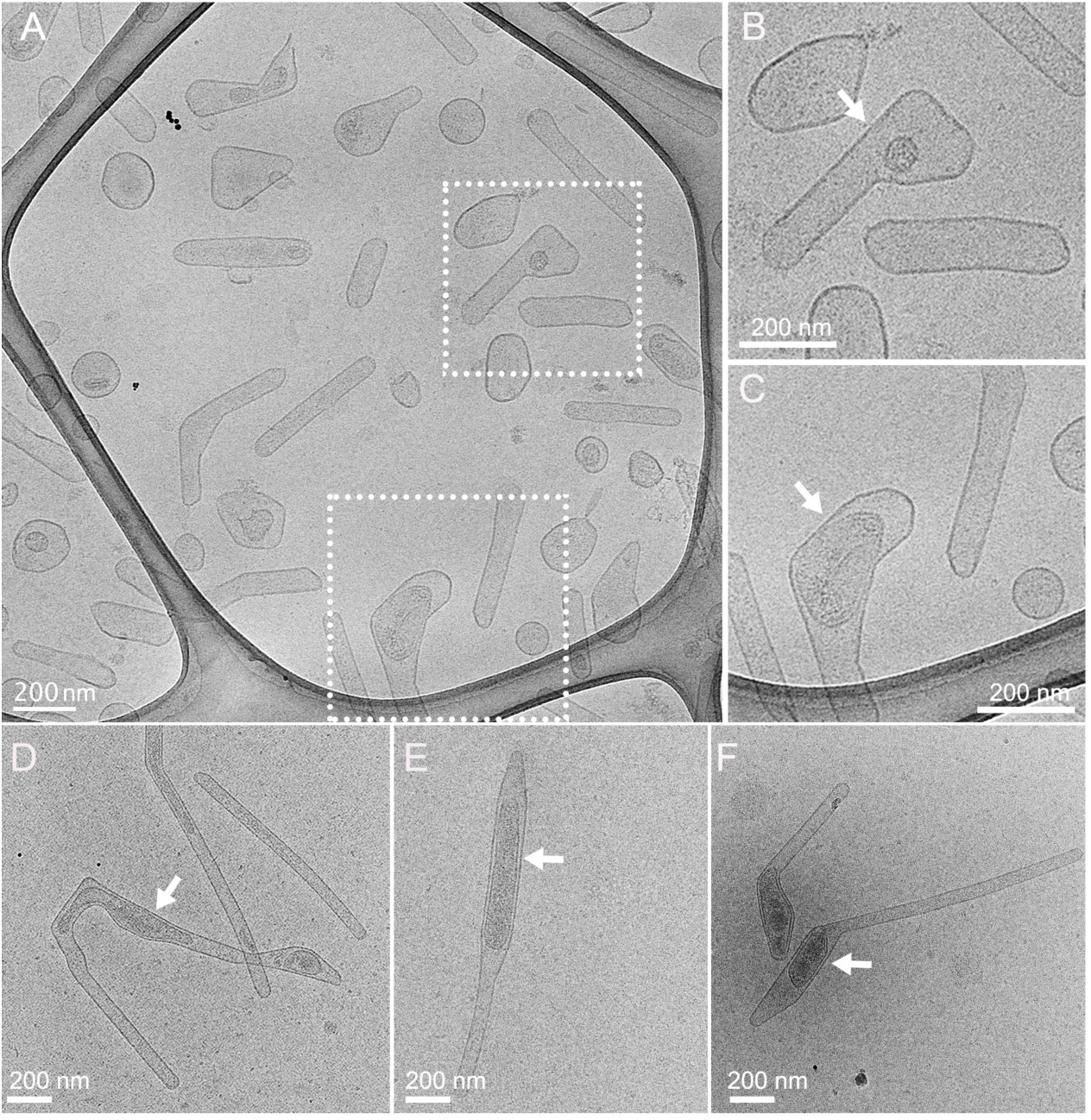
Cryo-EM imaging of vesicles released by *F. novicida* into the surrounding medium. Representative cryo-EM images of vesicles released by *F. novicida* U112 grown on BHI agar without cysteine supplementation. Panels B and C show enlarged views of the boxed regions in A. The arrows point to internal structures within the released OMT.

### Progression of *F. novicida* tube formation upon exposure to amino acid starvation

To investigate the progression of tube formation by *F. novicida*, we developed an assay to synchronize initiation of the bacterial response to amino acid starvation. *F. novicida* was grown in BHI broth to early log phase (OD_600_ ∼0.4), at which stage the cysteine in the medium was not yet depleted and OMT production was not induced^23^. The bacteria were then harvested and resuspended in cell-free, conditioned BHI medium obtained from stationary phase *F. novicida* cultures. This conditioned medium was depleted for cysteine, resulting in simultaneous exposure of the bacteria to amino acid starvation upon resuspension. We used both negative-stain, whole-bacteria transmission electron microscopy (TEM) and cryo-EM to examine the bacteria at time points just prior to resuspension in the conditioned BHI and at 1 and 6 h post resuspension.

Bacteria examined immediately prior to resuspension in the conditioned BHI were uniform in shape and lacked OMT (Fig. 4A, D, and G). The periplasm of these bacteria was mostly thin and evenly spaced, although some expanded regions were visible (Fig. 4G). Overall, the appearance of these bacteria was similar to the bacteria grown on cysteine-supplemented BHI agar (Fig. 1), consistent with growth in both cases under non-tube-inducing conditions. By 1 h post resuspension in the conditioned medium, tube formation had initiated on a subset of the bacteria and released OMT were visible on the EM grid (Fig. 4B and E). By cryo-EM, extended regions of the bacterial OM were present at the cell poles, with a corresponding expansion of the periplasm at these sites of protrusion (Fig. 4H). The OM extensions at the 1 h time point were broad at the base and with a rounded tip at the distal end, reflecting an early stage of tube formation. By 6 h post resuspension, there was a marked increase in tube-producing bacteria and released OMT compared to the 1 h time point (Fig. 4C and F). The OM projections were elongated and had obtained a more defined, tubular shape (Fig. 4I). Taken together, these results show that tube production by *F. novicida* is an early response following the transition of *F. novicida* from a nutrient-rich to a nutrient-depleted environment and involves a progressive restructuring of the OM, together with corresponding expansion of the periplasm and changes to the IM and cytoplasm at the base of the tubes.

**Figure 4.**
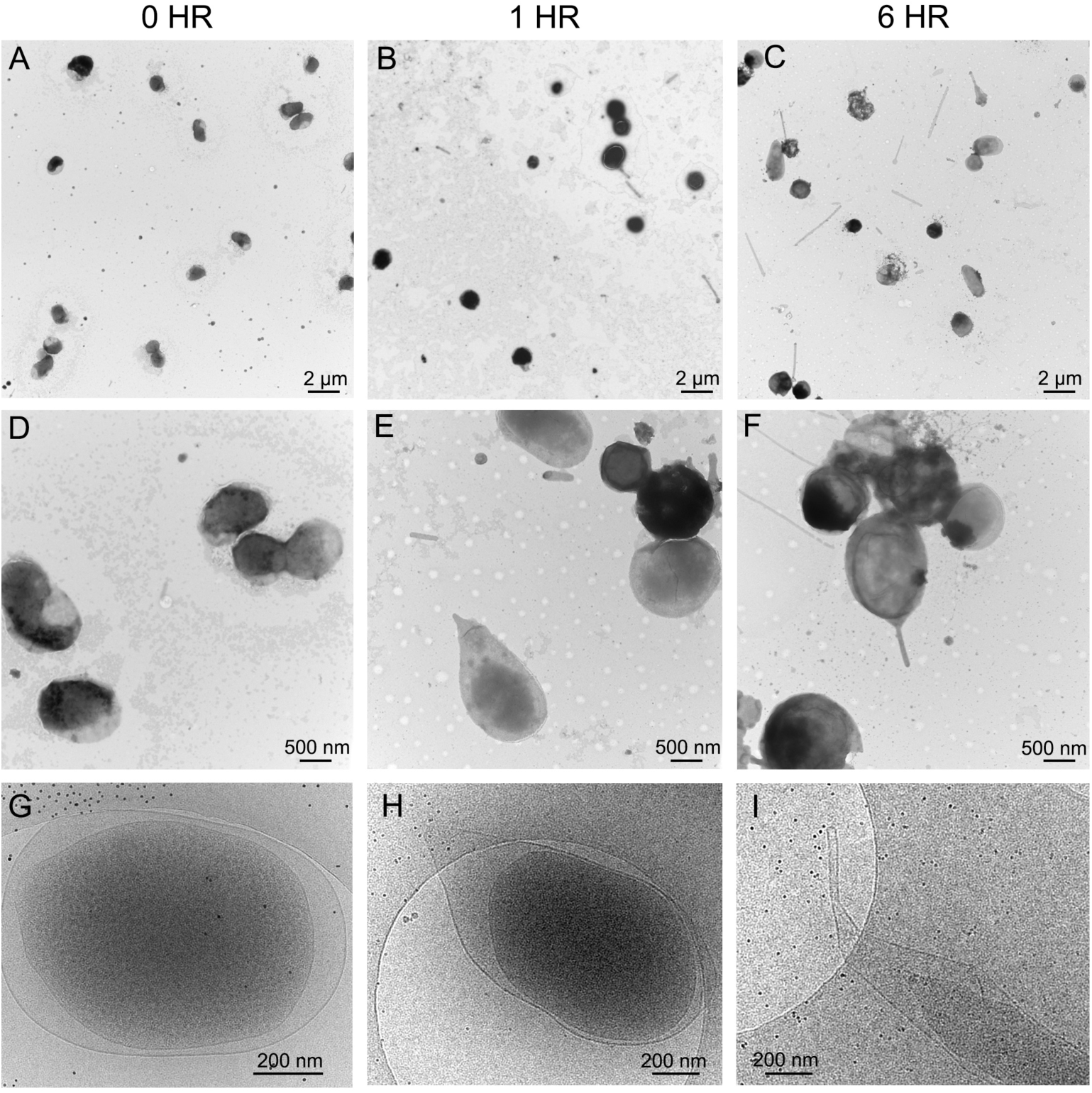
Progression of OMT formation upon exposure of *F. novicida* to amino acid starvation. Representative TEM (A-F) and cryo-EM (G-I) images of *F. novicida* U112 grown to early log phase in BHI medium and then resuspended into conditioned BHI medium depleted of cysteine. Bacteria were captured just prior to resuspension in the conditioned medium (0 h time point, panels A, D, and G), 1 h after resuspension (B, E, and H), and 6 h after resuspension (C, F, and I). Bacteria at 0 h show a regular morphology, with OMT formation increasing from 1 to 6 h.

### *F. novicida* tubes are produced during macrophage infection and colocalize with alterations in the phagosomal membrane

Previous studies showed that *Francisella* produces OMV and OMT during contact with macrophages and at early stages of infection, when the bacteria are still contained within the macrophage phagosome^10,23,30^. To investigate further the production of OMT during host cell infection, RAW 264.7 macrophage-like cells were infected with *F. novicida* U112 at a multiplicity of infection (MOI) of 2000, and at 20 min post-infection the infected macrophages were processed for thin-section, negative-stain TEM (Fig. 5A-C). The high MOI was used to aid detection of the bacteria in the sections and the early time point was chosen to capture bacteria at the macrophage surface or in the phagosomal compartment, prior to escape into the host cell cytosol. The TEM images confirmed the presence of OM protrusions for *F. novicida* that were interacting with pseudopod loops on the macrophage surface or located within phagosomes (Fig. 5B-C). The bacteria were irregular in shape and contained distended periplasmic regions, consistent with cultured bacteria grown under tube-inducing conditions (Figs. 2 and 4).

**Figure 5.**
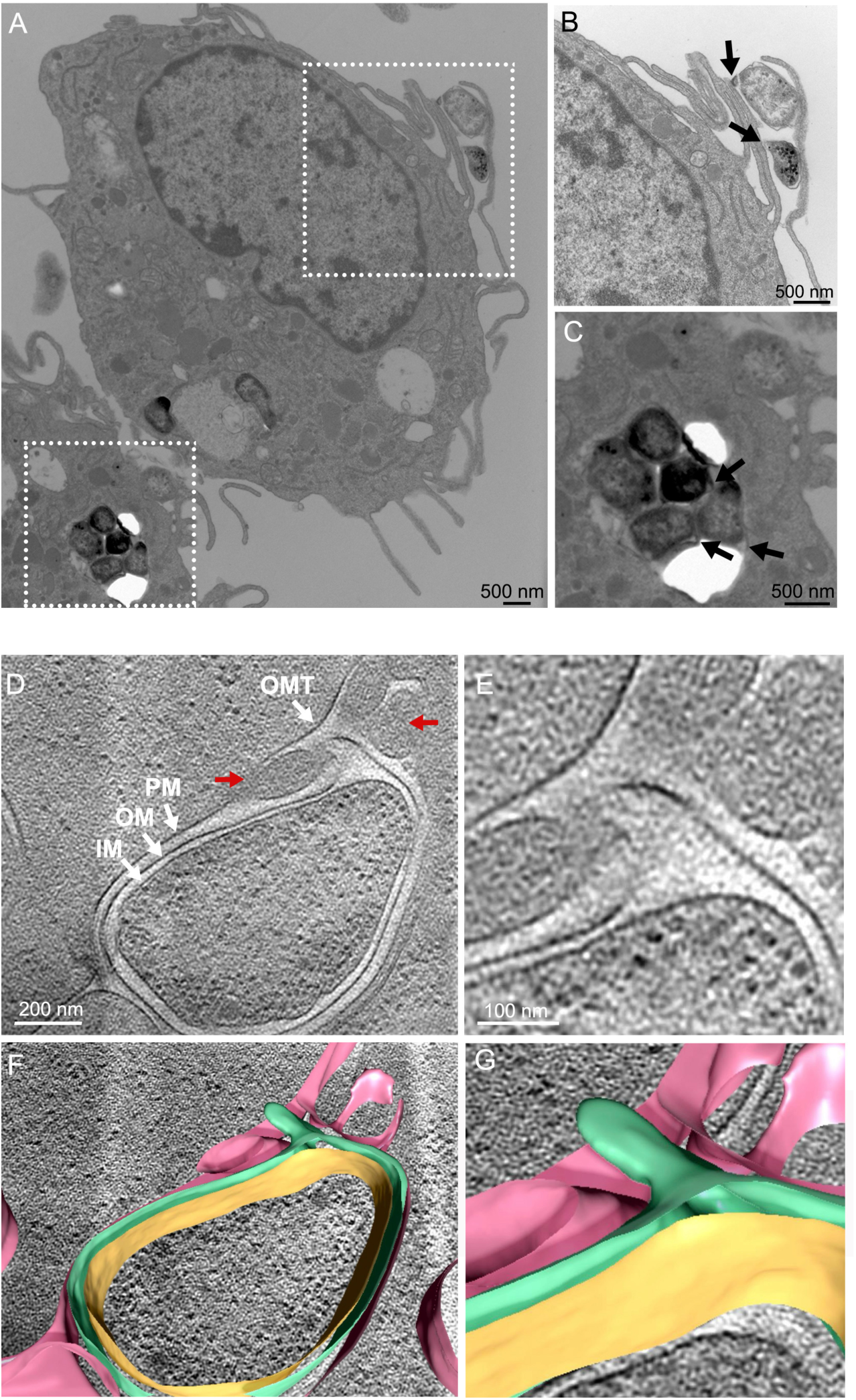
*F. novicida* produces OMT during macrophage infection and triggers phagosomal membrane rearrangement. (A-C) Thin-section TEM image of RAW 264.7 macrophage-like cells at 20 min post infection with *F. novicida* U112. Panels B and C are enlarged views of the boxed regions in panel A. The arrows indicate regions of OMT formation on bacteria at the macrophage cell surface (B) or within the phagosome (C). (D-G) Single section of a tomogram capturing an intracellular *F. novicida* in the macrophage phagosome. In panel D, the white arrows indicate the bacterial IM, OM, and OMT, and the host cell phagosomal membrane (PM). The red arrows indicate regions surrounding the OMT where the phagosomal membrane is undergoing vesiculation. Panel E is a zoomed-in image of the OMT region in panel D. (F and G) Overlaid tomogram section and 3D segmented images representing the bacterial IM (orange) and OM (green), and the host cell phagosomal membrane (pink).

As the OMT are difficult to capture by thin-section imaging and negative staining can alter cellular features, we next used cryo-FIB-ET to obtain a higher-resolution view of *F. novicida* within the RAW 264.7 phagosome in a near-native state. As shown in Fig. 5D-G, Movie S6, we were able to capture *F. novicida* enclosed within the macrophage phagosome. In the cryo-FIB-ET images and corresponding 3D segmentation analysis (Fig. 5F and G, Movie S6), the bacterial cytoplasm, IM, periplasm and OM were clearly defined, and the phagosomal membrane surrounding the bacterium was also clearly visible. Tubular extensions of the OM can be seen emerging from one pole of the bacteria, enclosing an enlarged periplasmic region (Fig. 5D-G, Movie S6). Notably, at the position where the tip of the tubular projection extends toward the phagosomal membrane, alterations of the phagosomal membrane were present, with the membrane appearing to vesiculate or break into smaller structures. This alteration may represent a step in the process of bacterial escape from the phagosome into the cytoplasm.

### The *F. novicida* T6SS colocalizes with OMT

The *Francisella* T6SS is essential for bacterial escape from the phagosome during intracellular infection. Given the observed changes to the phagosomal membrane that coincided with the presence of a tubular projection, we next investigated if the T6SS might localize to the OMT. To visualize the T6SS, we used a *F. novicida* U112 strain expressing a previously characterized fusion of IglA with sfGFP^18^. IglA forms the outer sheath of the T6SS structure that assembles in the bacterial cytoplasm. We grew the *iglA-sfGFP* stain on BHI agar under tube-inducing conditions (without cysteine supplementation) for 44-48 h and single colonies were picked for visualization by confocal microscopy. By brightfield imaging, the bacteria appeared pleomorphic, as expected for bacteria undergoing tube production, and OMT were observed extending from the bacterial surface and released into the surrounding medium (Fig. 6A). By fluorescence imaging, single, polarly localized IglA puncta were visible in the bacteria. Notably, where both IglA puncta and OMT were visible in the same bacterium, the puncta always localized at the base of where the tubes extended from the bacterial surface (Fig. 6A). In addition, we observed IglA puncta within OMT that were released from the bacteria (Fig. 6A), indicating the T6SS component is exported away from the bacteria with the tubular vesicles. We also examined bacteria grown under non-tube-inducing conditions (BHI agar supplemented with 0.1% cysteine). Under these conditions, the bacteria appeared more uniform in shape and OMT were not observed (Fig. 6B). However, single, polarly localized IglA puncta were still present in the bacteria. Taken together, our findings reveal that the *F. novicida* T6SS localizes to sites of OMT formation and remains associated with the OMT during their maturation and release from the bacteria.

**Figure 6.**
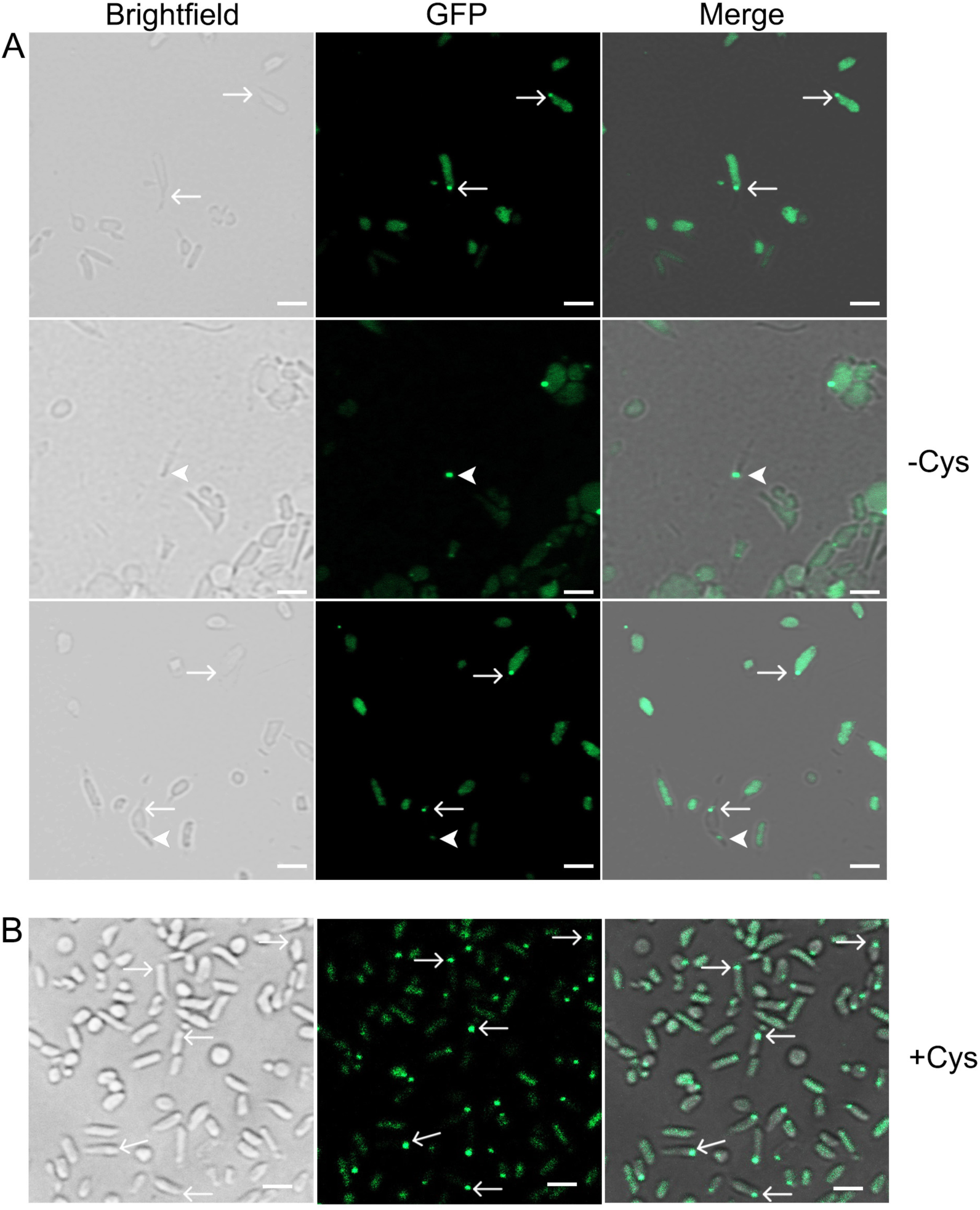
*F. novicida* T6SS colocalizes with OMT present on bacteria and released into the surrounding medium. (A) *F. novicida* U112 *iglA-sfGFP*, expressing the T6SS sheath protein IglA fused to GFP, was grown on BHI agar without cysteine supplementation (to induce OMT formation) and imaged by confocal microscopy. Brightfield, GFP, and merged images of the bacteria are shown. The arrows point to IglA puncta located at the base of OMT extending from the bacterial surface. The arrowheads point to IglA puncta present in OMT released into the surrounding medium. (B) *F. novicida* U112 *iglA-sfGFP* was grown on BHI agar with 0.1% cysteine supplementation (to repress OMT formation). The arrows point to IglA puncta located at a bacterial pole. Scale bars, 2 µm.

## DISCUSSION

Mechanisms governing the production and regulation of bacterial extracellular vesicles are not well understood. In addition to typical OMV, *Francisella* generates unique tubular extensions of its OM and releases tube-shaped vesicles into the surrounding medium^23^. The structural basis for formation of the *Francisella* OMT and their potential function during infection are unknown. In this study, we characterized the OMT ultrastructure and the steps leading to OMT generation in *F. novicida*. Our results revealed that major alterations in the bacterial envelope occur during tubulation and identified a dynamic cytoplasmic structure associated with OMT formation. Further, cryo-FIB-ET analysis of *F. novicida* during macrophage infection showed that OMT are produced within the macrophage phagosome and identified a potential role for the OMT in breakdown of the phagosomal membrane. This role was supported by our finding that the *F. novicida* T6SS, which is required for phagosomal escape, localized to the base of the tubular projections and was present in OMT released from the bacterial surface.

Typical extracellular vesicles produced by growing Gram-negative bacteria bud off from the OM and contain OM and periplasmic cargo^26–29^. The *F. novicida* OMT are also formed from the OM^23^. However, the cryo-ET visualization in this study revealed that the OMT include tubular extensions of the IM and are associated with a cytoplasmic structure that itself undergoes changes in shape and extends partly into the growing tube. Therefore, the OMT are derived from both the OM and IM and contain components from both membranes, along with periplasmic and cytoplasmic material enclosed by the membranes. We observed this composition for OMT extending from the bacterial surface as well as released extracellularly. Consistent with this, proteomic analysis of mixed OMV and OMT purified from *F. novicida* reported that cytoplasmic and IM proteins comprised ∼40% of the total abundance of the constituent proteins^23^. A variety of tubular membrane extensions have been reported for Gram-negative bacteria^34,37–40^. The *F. novicida* OMT appear distinct from these other reported structures, not only in their shape, dynamic nature, and regulation, but also for the involvement of a cytoplasmic organelle in tubulation and the apparent origin of the OMT at the bacterial IM.

Using a synchronized OMT-induction assay, we detected progressive changes in the *F. novicida* cell envelope during tube initiation and maturation that occurred over a 6 h period (Fig. 4). Thus, OMT formation is driven by a defined program that is responsive to environmental cues (amino acid starvation). *F. novicida* grown in amino acid sufficient conditions were mostly regular in shape but still exhibited notable differences in their envelope compared to canonical Gram-negative bacteria such as *E. coli*, including enlarged regions of the periplasm at the cell poles and uneven sections of the IM and OM bilayers that were sometimes associated with periplasmic densities (Fig. 1). These features raise intriguing questions regarding the structural organization of the cell envelope in *Francisella* spp., such as the placement of the peptidoglycan cell wall and connections among the cell wall, IM and OM. In addition to the unusual architecture of the cell envelope, we also observed an internal cytoplasmic structure that was consistently associated with OMT formation and present under both amino acid sufficient and depleted conditions. The cytoplasmic structure had the appearance of a membrane-bound organelle, with internal contents separated from the surrounding cytoplasm, although its nature and composition remain to be determined. Under amino acid sufficient conditions, the cytoplasmic structure was generally oval shaped and located at the cell poles (Fig. 1). The structure exhibited notable changes from an oval shape under cysteine sufficient conditions to a more bulb-like conformation that extended into the tubular projection, along with cytoplasmic material, as OMT formation progressed (Fig. 2). Based on its positioning at the base of the tubes and its progressive changes during OMT formation, we hypothesize that the cytoplasmic structure serves as a specialized apparatus that controls tubulation by initiating extension of the IM, which then exerts outward pressure on the OM to drive OMT formation. At later stages of OMT formation, the tubular projections became narrower, the periplasmic space more constricted, and the cytoplasmic structure assumed a fragmented appearance. These changes likely reflect steps leading to breakdown of the tubulation machinery in preparation for OMT release into the extracellular environment.

We previously observed the production of OMT by *F. novicida* during the initial stages of host cell infection, upon contact with the macrophage cell surface and within the phagosome^23^. This phenomenon is not restricted to *F. novicida*, as *F. tularensis* subsp. *holarctica* produces tubular protrusions and vesicles during early stages of macrophage infection^10,30^ and *F. tularensis* subsp. *tularensis* also produces OMT^25,41^. Here, using cryo-FIB-ET, we captured high-resolution images of *F. novicida* contained within the macrophage phagosome (Fig. 5). These images verified that the bacteria produce OMT during infection. Notably, where the OMT extended toward the phagosomal membrane, the phagosomal membrane appeared to fragment into bleb-like structures surrounding the tip of the bacterial tube. Previous studies, using thin-section TEM, also noted vesiculation and fragmentation of the phagosomal membrane following infection of macrophages with *F. tularensis*^12,30^. Rapid escape from the phagosomal compartment is a hallmark of *Francisella* pathogenesis^1–3^. In line with these studies and our current findings, we propose that the observed perturbation of the phagosomal membrane represents a step in bacterial escape from the phagosome into the cytoplasm.

Essential for *Francisella’s* escape from the phagosome is the T6SS^14–18^. The T6SS is a contractile puncturing device, related to phage tail complexes^42,43^. Upon activation, the T6SS injects effector proteins directly into neighboring bacterial or eukaryotic cells, promoting bacterial survival and proliferation. By examining *F. novicida* expressing GFP-tagged IglA, which forms the outer sheath of the contractile structure that assembles in the bacterial cytoplasm, we found that the *Francisella* T6SS co-localizes with the OMT (Fig. 6). IglA was observed both at the base of tubes extending from the bacterial surface, as well as contained within OMT released outside the bacteria. In agreement with this, T6SS components have been detected by proteomics analysis of OMV and OMT purified from *Francisella* spp.^23–25,41^. Future studies are needed to examine T6SS activity and dynamics in the context of the OMT and host cell infection; nevertheless, the co-localization between the T6SS, which is required for phagosomal escape, and the OMT, which was found at regions of phagosomal disruption, suggests a functional association between these two elements. In contrast to other bacteria, where T6SSs are preferentially assembled along the long (narrow) axis of the cell, the *Francisella* T6SS assembles at the cell poles^18^. This distinct positioning is consistent with the polar localization observed for the OMT and reinforces the relationship between these structures. Further, both the T6SS and OMT are regulated by amino acid starvation, a condition encountered by the bacteria within the macrophage phagosome^25,33,44,45^. The *F. novicida* T6SS localization pattern also mirrors that of the internal cytoplasmic structure that we found to be associated with OMT formation. Like the cytoplasmic structure, the T6SS puncta were observed in bacteria grown under both OMT noninducing and inducing conditions, indicating that the presence of the T6SS is not sufficient to drive membrane tubulation. These results, along with prior and ongoing analyses^25^, suggest that the T6SS is not required for formation of the OMT. Rather, we hypothesize that the T6SS in *Francisella* may have evolved to localize with and take advantage of the OMT to facilitate effector delivery to the host during infection. The OMT extending from the bacterial surface may facilitate contact of the T6SS with the phagosomal membrane, and OMT released from the bacteria may extend the reach of the T6SS away from the bacterial surface.

In conclusion, our findings demonstrate that OMT production by *Francisella* is a regulated process involving a gradual and progressive remodeling of the bacterial cell envelope under amino acid-starved conditions. Furthermore, we identify a novel cytoplasmic structure that exhibits remarkable shape alterations in tube-forming bacteria and may drive tube formation, initiating at the bacterial IM. Our observations reveal the association of OMT produced by *F. novicida* contained within the macrophage phagosome with fragmented regions of the phagosomal membrane. We identified a convergence between the *Francisella* OMT and the T6SS, suggesting that the bacterial tubes and released OMT may facilitate the delivery of T6SS effectors to the phagosomal membrane, enabling bacterial escape to the cytoplasm. Although the precise role of the *Francisella* OMT in phagosomal escape and subsequent virulence requires further investigation, our study lays a foundation for understanding the molecular mechanism of bacterial tubulation and its role during *Francisella* pathogenesis.

## MATERIALS AND METHODS

### Bacterial strains and growth conditions

*F. novicida* strains U112 (BEI Resources, ATCC 15482) and U112 *iglA-sfGFP*^18^ were grown in BHI (brain heart infusion) medium [37 g/l BHI powder (BD Biosciences), adjusted to pH 6.8] or BHI supplemented with 0.1% cysteine. For plates, Bacto agar (BD Biosciences) was added to BHI medium at 15 g/l. Bacteria grown on plates were incubated at 37°C in the presence of 5% CO_2_. For liquid cultures, the media was incubated in the presence of 5% CO_2_ for 30 min prior to bacterial inoculation. Liquid cultures were grown at 37°C with aeration by shaking at 100 rpm. Liquid cultures were started from frozen stocks by direct inoculation, followed by overnight growth. The cultures were then diluted 1:100 to achieve an optical density at 600 nm (OD_600_) of approximately 0.01. Day cultures were grown until exponential phase (OD_600_ of 0.5–0.8) or stationary phase (OD_600_ of 1.2–1.4).

### Purification of vesicles released by *F. novicida*

*F. novicida* U112 lawns from four BHI agar plates (without 0.1% cysteine supplementation), grown for 44-48 h, were scraped and suspended in 7.5 ml of OMV Buffer (20 mM HEPES, pH 7.5, 150 mM NaCl, 0.05% sodium azide). The resuspended bacteria were centrifuged at 9780 x g for 10 min at 4°C. The supernatant containing vesicles was carefully collected and transferred into a fresh tube. The bacterial pellet was washed twice with 20 ml of OMV Buffer and the wash was added to the transferred supernatant fraction. The collected supernatants were pooled and filtered through a 0.2 µm PES membrane (Millipore). The filtered supernatants were then subjected to ultracentrifugation at 100,000 x g for 1 h at 4°C. The resulting vesicle pellet was resuspended in 500 µl of OMV Buffer and stored at −20°C until analysis by cryo-EM.

### Cryo-EM of vesicles released by *F. novicida*

A 4 μl aliquot of the purified vesicles was applied to a freshly glow-discharged (Edwards) lacey carbon grid covered with a thin layer of continuous carbon film. After one minute incubation, excess solution on the grid was blotted with a piece of filter paper and then the grid was rapidly plunged into liquid ethane using a Vitrobot (FEI). Low dose imaging (15 e/Å) was performed in a JEM 2010F transmission electron microscope (JEOL USA) operating at high tension of 200 kV and magnification of 50,000, with an objective lens under-focus value of ∼3-5 μm and with the EM grids maintained at −170°C in a Gatan 626 cryo-specimen holder. Digital micrographs were recorded on a Gatan 4K by 4K UltraScan CCD camera.

### OMT induction assay

To generate conditioned BHI medium depleted for cysteine, overnight cultures of *F. novicida* were grown in BHI to early stationary phase (OD_600_ 1.0–1.2, ∼10 h). The bacteria were then pelleted by centrifugation at 9780 x g for 10 min at 4°C. The resulting supernatants were collected and filtered through a 0.2 µm PES membrane (Millipore) to obtain the cell-free conditioned BHI medium. Separately, day cultures of *F. novicida* were grown in 25 ml standard BHI (without cysteine supplementation) to early log phase (OD_600_ of 0.3–0.4, ∼3 h). Bacteria were pelleted at 9780 x g for 10 min at 4°C and resuspended in 25 ml of previously prepared conditioned BHI, pre-warmed for 30 min at 37°C with 5% CO_2_. The cultures were further incubated in conditioned BHI for 6 h. The bacteria were analyzed by TEM or cryo-ET just before resuspension in the conditioned medium and at 1 and 6 h post resuspension in the conditioned medium.

### TEM analysis of whole bacteria

For analysis of whole bacteria grown in liquid culture for the OMT induction assay, 1 ml of culture was centrifuged (9780 x g, 5 min, 4°C) and resuspended in 200 µl sterile PBS. Samples were placed on polyvinyl formvar-coated copper grids (Electron Microscopy Services) and allowed to adhere for 2 min. Subsequently, the grids were treated with 1% glutaraldehyde for 1 min, washed twice with PBS, twice with water, stained with 0.5% phosphotungstic acid for 20 s, and air dried. Grids for TEM were examined using an FEI Tecnai12 BioTwinG2 electron microscope operating at an accelerating voltage of 80 kV. Images were captured using an AMT XR-60 charge-coupled device digital camera system.

### Preparation of RAW 264.7 macrophage-like cells

RAW 264.7 macrophage-like cells (ATCC TIB-71) from a cryogenic vial were thawed at 37°C in a water bath. Thawed macrophages were transferred to a 15 ml conical tube and centrifuged at 100 x g for 5 min at room temperature. After discarding the supernatant, the pellet was resuspended in 7 ml of pre-warmed DMEM (Dulbecco’s modified Eagle’s medium) with 10% FBS (fetal bovine serum) and transferred to a T-25 flask. Macrophages were incubated at 37°C with 5% CO_2_ and passaged twice to ensure they were actively dividing.

### *F. novicida* infection of RAW 264.7 cells for TEM analysis

RAW 264.7 cells were passaged 18-24 h prior to infection to ensure 90% cell viability and 80% cell confluency in the T-25 flask at the time of infection. Macrophages were removed from the T-25 flask and resuspended in DMEM with 10% FBS (fetal bovine serum) to a final concentration of 6 x 10^6^. The macrophages were pelleted (1000 x g for 10 min at 4°C) and the supernatant was discarded. Day cultures of *F. novicida* were grown in BHI to early log phase (OD_600_ = 0.4) and 1.2 x 10^10^ bacterial cells were centrifuged (9780 x g for 10 min at 4°C) and resuspended in 1 ml DMEM with 10% FBS. The resuspended bacteria were added to the pelleted macrophages at a MOI of 2000. The cells were centrifuged twice (200 x g and 800 x g, 10 min each at 4°C) to facilitate contact and supernatants were discarded. The tube containing the pelleted bacteria and macrophages was incubated in a 37°C water bath for 20 min to initiate phagocytosis. The bacteria and macrophages were then treated with 1 ml of 2.5% glutaraldehyde at 37°C for 2 min, followed by incubation on ice for 30 min. The cells were centrifuged at 10,000 × g for 10 min at 4°C, and the resulting pellet was resuspended in 1 ml of ice-cold phosphate-buffered saline (PBS). For thin section analysis, samples were initially fixed in 2.5% EM grade glutaraldehyde in 0.1 M PBS, pH 7.4, for a minimum of 1 h. Subsequently, the samples were treated with 1% osmium tetroxide in 0.1 M PBS, dehydrated through a series of graded ethyl alcohol solutions, and embedded in Durcupan resin. Thin sections measuring 80 nm in thickness were sliced using a Reichert-Jung UltracutE ultramicrotome and placed onto formvar-coated slot copper grids. These sections were then stained with uranyl acetate and lead citrate. Grids for TEM were examined using an FEI Tecnai12 BioTwinG2 electron microscope operating at an accelerating voltage of 80 kV. Images were captured using an AMT XR-60 charge-coupled device digital camera system.

### Bacterial sample preparation for Cryo-ET

*F. novicida* U112 was grown under cysteine sufficient and OMT-inducing conditions as described above. The bacteria were pelleted by centrifugation at 9,780 x g for 10 min at 4°C, resuspended in PBS, and adjusted to an OD_600_ of 1.0. The bacterial suspension was mixed with BSA-coated 10 nm Gold Tracer beads (Aurion), and 5 μl of the mixture was deposited onto glow-discharged cryo-EM grids (Quantifoil, Cu, 200 mesh, R2/1). The grids were set on a homemade gravity plunger and were blotted at the back with filter paper for almost 4 sec. Then, the grids were immediately plunge-frozen in liquid ethane.

### *F. novicida* infection of RAW 264.7 cells for cryo-focused ion beam (cryo-FIB) milling

RAW 246.7 cells were prepared as described above. After passage to a new flask, the cells were grown overnight on cryo-EM grids (Quantifoil, Au, 200 mesh, R1/4). Glow-discharged cryo-EM grids were treated with poly-D-lysine and washed by DMEM several times before seeding cells on the grids. For bacterial infection, *F. novicida* U112 was grown in BHI medium overnight. The overnight culture was then inoculated and grown in fresh BHI medium until it reached an OD_600_ of 0.4. A 3 ml aliquot of the bacterial culture was pelleted by centrifugation at 2000 x g for 5 min and resuspended in DMEM. Then, the RAW 246.7 culture medium was removed and replaced with the bacterial suspension in DMEM. To initiate infection on the cryo-EM grids, co-culture samples were centrifuged at 800 x g for 10 min and incubated at 37°C with 5.0% CO_2_ for 20 min. The DMEM was replaced with DMEM + 10% glycerol, and the cryo-EM grids were then set on a homemade plunger to prepare frozen specimens for cryo-focused ion beam (cryo-FIB) milling.

### Cryo-focused ion beam (cryo-FIB) sample preparation

The frozen specimens of RAW 246.7 cells infected with *F. novicida* U112 were loaded into an Aquilos cryo FIB-SEM system (Thermo Fisher Scientific) maintained at around −180°C. To protect specimens from the Gallium ion beam, organic and inorganic platinum layers were coated on the entire cryo-EM grids. After finding targets using ion and electron beam images, the top parts of the RAW 246.7 cells were milled by the ion beam at 0.3 nA current. Once intracellular bacteria were visualized in SEM images, the bottom parts of the RAW 246.7 cells were milled. At ∼1.0 μm sample thickness, the ion beam current was reduced to 0.1 nA and used to mill target cells to ∼0.5 μm thickness. In the last step, the ion beam current was further reduced to 50 and 30 pA and used to polish lamellae to a final thickness of ∼150 nm. The milling angle was 8° for all target cells. After all lamellae were prepared, another inorganic platinum layer was coated on their surface using a shorter sputtering time.

### Cryo-ET data collection and processing

The frozen specimens of *F. novicida* U112 were recorded at almost −180°C using a Titan Krios G2 300 kV transmission electron microscope (Thermo Fisher Scientific) equipped with a field emission gun, K2 detector, and a BioQuantum imaging filter (Gatan). To operate the microscope, SerialEM software^46^ was used for automated acquisition of tilt series images at a magnification with a physical pixel size of 5.46 Å. The microscope stage was tilted from −45° to +45° in 3° increments using the dose-symmetric scheme in a SerialEM tilt series program. During data acquisition for cryo-FIB samples, image recording began at 8° or −8° stage angle, and tilt series images were collected from ±40° to ±56° in 3° increments using the dose-symmetric scheme in FastTomo script^47^. The positive or negative tilt angles depended on the direction of lamellae in the microscope. A K3 detector (Gatan) was used to record images from the cryo-FIB samples. Motioncor2^48^ was used to correct electron beam-induced image drifts. Then, IMOD software was used to create image stacks and align images in each tilt series by tracking fiducial gold beads^49,50^. For images from cryo-FIB samples, small dots from the last sputtering step in cryo-FIB milling were used to align images in each tilt series. 4x binned images were generated by binvol command in IMOD, and then 4× binned tomograms with simultaneous iterative reconstruction technique (SIRT) were reconstructed using Tomo3D^51^. 4× binned tomograms were imported to Dragonfly software (version 2022.2, Comet Technologies Canada) for 3D visualization, and then bacterial OM, IM, and oval-shaped cytoplasmic structures were segmented and pseudo colored. For ribosomes in the images, round-shaped features in the software were mapped back into the 3D images.

### Fluorescence microscopy

*F. novicida* U112 *iglA-sfGFP* was grown on BHI agar for 44–48 h, with or without supplementation with 0.1% cysteine. Approximately 15-20 bacterial colonies were scraped off, resuspended together in 1 ml sterile PBS, and centrifuged at 9780 x g for 3 min to harvest the cells. The supernatant was discarded, and the remaining cell pellet was resuspended in 500 µl PBS. Approximately 5–8 µl of resuspended cells were placed on a pad of 1% agarose in sterile PBS, followed by a single drop of SlowFade Gold antifade reagent (Invitrogen), and sealed with a coverslip. Samples were imaged immediately using the Zeiss LSM 980 Airyscan 2 NLO Two-Photon Confocal Microscope.

## ACKNOWLEDGEMENTS

We thank Susan van Horn and Guowei Tian of the Stony Brook University Central Microscopy Imaging Center for assistance with the TEM and confocal experiments, respectively. We thank Marek Basler (University of Basel) for providing the U112 *iglA-sfGFP* strain. We thank Vinaya Sampath (Long Island University), Marek Basler, and Maj Brodmann (Rockefeller University) for helpful discussions.

This work was supported by National Institute of Allergy and Infectious Diseases (NIAID), National Institutes of Health (NIH) award R01 AI141633 (D.G.T) and a Stony Brook University-Brookhaven National Laboratory Seed Grant (D.G.T.). S.T., S.Z., H.Z., and J.L. were supported by grants R01 AI152421, R01 AI087946 and R01 AI132818 from the NIAID. Cryo-ET data were collected at the Yale CryoEM resource, which is funded in part by NIH grant 1S10OD023603-01A1.

## SUPPLEMENTARY INFORMATION

**Movie S1. Cryo-ET of *F. novicida* U112 grown on BHI agar supplemented with 0.1% cysteine supplementation (OMT-repressing condition)**. Corresponds to the tomogram slice shown in Fig. 1A.

**Movie S2. Cryo-ET of *F. novicida* U112 grown on BHI agar supplemented with 0.1% cysteine supplementation (OMT-repressing condition)**. Corresponds to the tomogram slice shown in Fig. 1B.

**Movie S3. Cryo-ET of *F. novicida* U112 grown on BHI agar supplemented without cysteine supplementation (OMT-inducing condition)**. Corresponds to the tomogram slice shown in Fig. 2A.

**Movie S4. Cryo-ET of *F. novicida* U112 grown on BHI agar supplemented without cysteine supplementation (OMT-inducing condition)**. Corresponds to the tomogram slice shown in Fig. 2B.

**Movie S5. Cryo-ET of *F. novicida* U112 grown on BHI agar supplemented without cysteine supplementation (OMT-inducing condition)**. Corresponds to the tomogram slice shown in Fig. 2D.

**Movie S6. Cryo-ET of *F. novicida* U112 within the phagosome of a RAW 264.7 cell.** The tomogram is overlaid with a 3D segmentation analysis showing the bacterial IM (orange) and OM (green), and the host cell phagosomal membrane (pink). Corresponds to the tomogram slice shown in Fig. 5D.

## Notes

### Competing Interest Statement

The authors have declared no competing interest.

## REFERENCES

1. Oyston, P.C.F. *Francisella tularensis*: unravelling the secrets of an intracellular pathogen. J Med Microbiol 57, 921–930 (2008).

2. Celli, J. & Zahrt, T.C. Mechanisms of *Francisella tularensis* intracellular pathogenesis. Cold Spring Harb Perspect Med 3, a010314 (2013).

3. Steiner, D.J., Furuya, Y. & Metzger, D.W. Host-pathogen interactions and immune evasion strategies in *Francisella tularensis* pathogenicity. Infect Drug Resist 7, 239–251 (2014).

4. Ellis, J., Oyston, P.C., Green, M. & Titball, R.W. Tularemia. Clin Microbiol Rev 15, 631–646 (2002).

5. McCrumb, F.R. Aerosol infection of man with *Pasteurella tularensis*. Bacteriol Rev 25, 262–267 (1961).

6. Oyston, P.C., Sjostedt, A. & Titball, R.W. Tularaemia: bioterrorism defence renews interest in *Francisella tularensis*. Nat Rev Microbiol 2, 967–978 (2004).

7. Rohmer, L. et al. Comparison of *Francisella tularensis* genomes reveals evolutionary events associated with the emergence of human pathogenic strains. Genome Biol 8, R102 (2007).

8. Jones, C.L. et al. Subversion of host recognition and defense systems by *Francisella* spp. Microbiol Mol Biol Rev 76, 383–404 (2012).

9. Benziger, P.T., Kopping, E.J., McLaughlin, P.A. & Thanassi, D.G. *Francisella tularensis* disrupts TLR2-MYD88-p38 signaling early during infection to delay apoptosis of macrophages and promote virulence in the host. mBio 14, e0113623 (2023).

10. Golovliov, I., Baranov, V., Krocova, Z., Kovarova, H. & Sjostedt, A. An attenuated strain of the facultative intracellular bacterium *Francisella tularensis* can escape the phagosome of monocytic cells. Infect Immun 71, 5940–5950 (2003).

11. Anthony, L.D., Burke, R.D. & Nano, F.E. Growth of *Francisella* spp. in rodent macrophages. Infect Immun 59, 3291–3296 (1991).

12. Clemens, D.L., Lee, B.Y. & Horwitz, M.A. Virulent and avirulent strains of *Francisella tularensis* prevent acidification and maturation of their phagosomes and escape into the cytoplasm in human macrophages. Infect Immun 72, 3204–3217 (2004).

13. Checroun, C., Wehrly, T.D., Fischer, E.R., Hayes, S.F. & Celli, J. Autophagy-mediated reentry of *Francisella tularensis* into the endocytic compartment after cytoplasmic replication. Proc Natl Acad Sci U S A 103, 14578–14583 (2006).

14. Clemens, D.L., Ge, P., Lee, B.Y., Horwitz, M.A. & Zhou, Z.H. Atomic structure of T6SS reveals interlaced array essential to function. Cell 160, 940–951 (2015).

15. Barker, J.R. et al. The *Francisella tularensis* pathogenicity island encodes a secretion system that is required for phagosome escape and virulence. Mol Microbiol 74, 1459–1470 (2009).

16. Nano, F.E. et al. A *Francisella tularensis* pathogenicity island required for intramacrophage growth. J Bacteriol 186, 6430–6436 (2004).

17. Eshraghi, A. et al. Secreted effectors encoded within and outside of the *Francisella* Pathogenicity Island promote intramacrophage growth. Cell Host Microbe 20, 573–583 (2016).

18. Brodmann, M., Dreier, R.F., Broz, P. & Basler, M. *Francisella* requires dynamic type VI secretion system and ClpB to deliver effectors for phagosomal escape. Nat Commun 8, 15853 (2017).

19. Hager, A.J. et al. Type IV pili-mediated secretion modulates Francisella virulence. Mol Microbiol 62, 227–237 (2006).

20. Gil, H. et al. Deletion of TolC orthologs in *Francisella tularensis* identifies roles in multidrug resistance and virulence. Proc Natl Acad Sci U S A 103, 12897–12902 (2006).

21. Gil, H., Benach, J.L. & Thanassi, D.G. Presence of pili on the surface of *Francisella tularensis*. Infect Immun 72, 3042–3047 (2004).

22. Pierson, T. et al. Proteomic characterization and functional analysis of outer membrane vesicles of Francisella novicida suggests possible role in virulence and use as a vaccine. J Proteome Res 10, 954–967 (2011).

23. McCaig, W.D., Koller, A. & Thanassi, D.G. Production of outer membrane vesicles and outer membrane tubes by *Francisella novicida*. J Bacteriol 195, 1120–1132 (2013).

24. Klimentova, J. et al. Francisella tularensis subsp. holarctica Releases Differentially Loaded Outer Membrane Vesicles Under Various Stress Conditions. Front Microbiol 10, 2304 (2019).

25. Sampath, V., McCaig, W.D. & Thanassi, D.G. Amino acid deprivation and central carbon metabolism regulate the production of outer membrane vesicles and tubes by *Francisella*. Mol Microbiol 107, 523–541 (2018).

26. Ellis, T.N. & Kuehn, M.J. Virulence and immunomodulatory roles of bacterial outer membrane vesicles. Microbiol Mol Biol Rev 74, 81–94 (2010).

27. Schwechheimer, C. & Kuehn, M.J. Outer-membrane vesicles from Gram-negative bacteria: biogenesis and functions. Nat Rev Microbiol 13, 605–619 (2015).

28. Beveridge, T.J. Structures of Gram-negative cell walls and their derived membrane vesicles. J Bacteriol 181, 4725–4733 (1999).

29. Schertzer, J.W. & Whiteley, M. Bacterial outer membrane vesicles in trafficking, communication and the host-pathogen interaction. J Mol Microbiol Biotechnol 23, 118–130 (2013).

30. Pavkova, I. et al. Francisella tularensis Outer Membrane Vesicles Participate in the Early Phase of Interaction With Macrophages. Front Microbiol 12, 748706 (2021).

31. Lauriano, C.M. et al. MglA regulates transcription of virulence factors necessary for *Francisella tularensis* intraamoebae and intramacrophage survival. Proc Natl Acad Sci U S A 101, 4246–4249 (2004).

32. Guina, T. et al. MglA regulates *Francisella tularensis* subsp. *novicida* (*Francisella novicida*) response to starvation and oxidative stress. J Bacteriol 189, 6580–6586 (2007).

33. Charity, J.C., Blalock, L.T., Costante-Hamm, M.M., Kasper, D.L. & Dove, S.L. Small molecule control of virulence gene expression in *Francisella tularensis*. PLoS Pathog 5, e1000641 (2009).

34. Kaplan, M. et al. In situ imaging of bacterial outer membrane projections and associated protein complexes using electron cryo-tomography. Elife 10(2021).

35. Krypotou, E. et al. Bacteria require phase separation for fitness in the mammalian gut. Science 379, 1149–1156 (2023).

36. Protter, D.S.W. & Parker, R. Principles and Properties of Stress Granules. Trends Cell Biol 26, 668–679 (2016).

37. Galkina, S.I. et al. Membrane tubules attach Salmonella Typhimurium to eukaryotic cells and bacteria. FEMS Immunol Med Microbiol 61, 114–124 (2011).

38. Pirbadian, S. et al. Shewanella oneidensis MR-1 nanowires are outer membrane and periplasmic extensions of the extracellular electron transport components. Proc Natl Acad Sci U S A 111, 12883–12888 (2014).

39. Wei, X., Vassallo, C.N., Pathak, D.T. & Wall, D. Myxobacteria produce outer membrane-enclosed tubes in unstructured environments. J Bacteriol 196, 1807–1814 (2014).

40. Fischer, T. et al. Biopearling of Interconnected Outer Membrane Vesicle Chains by a Marine Flavobacterium. Appl Environ Microbiol 85(2019).

41. Klimentova, J. et al. Cross-species proteomic comparison of outer membrane vesicles and membranes of *Francisella tularensis* subsp. *tularensis* versus subsp. *holarctica*. J Proteome Res 20, 1716–1732 (2021).

42. Bingle, L.E., Bailey, C.M. & Pallen, M.J. Type VI secretion: a beginner’s guide. Curr Opin Microbiol 11, 3–8 (2008).

43. Cianfanelli, F.R., Monlezun, L. & Coulthurst, S.J. Aim, Load, Fire: The Type VI Secretion System, a Bacterial Nanoweapon. Trends Microbiol 24, 51–62 (2016).

44. Appelberg, R. Macrophage nutriprive antimicrobial mechanisms. J Leukoc Biol 79, 1117–1128 (2006).

45. Zhang, Y.J. & Rubin, E.J. Feast or famine: the host-pathogen battle over amino acids. Cell Microbiol 15, 1079–1087 (2013).

46. Mastronarde, D.N. Automated electron microscope tomography using robust prediction of specimen movements. J Struct Biol 152, 36–51 (2005).

47. Xu, A. & Xu, C. FastTomo: A SerialEM Script for Collecting Electron Tomography Data. bioRxiv, 2021.2003.2016.435675 (2021).

48. Zheng, S.Q. et al. MotionCor2: anisotropic correction of beam-induced motion for improved cryo-electron microscopy. Nat Methods 14, 331–332 (2017).

49. Kremer, J.R., Mastronarde, D.N. & McIntosh, J.R. Computer visualization of three-dimensional image data using IMOD. J Struct Biol 116, 71–76 (1996).

50. Mastronarde, D.N. & Held, S.R. Automated tilt series alignment and tomographic reconstruction in IMOD. J Struct Biol 197, 102–113 (2017).

51. Agulleiro, J.I. & Fernandez, J.J. Tomo3D 2.0--exploitation of advanced vector extensions (AVX) for 3D reconstruction. J Struct Biol 189, 147–152 (2015).

